# KDML: a machine-learning framework for inference of multi-scale gene functions from genetic perturbation screens

**DOI:** 10.1101/761106

**Authors:** Heba Z. Sailem, Jens Rittscher, Lucas Pelkmans

## Abstract

Characterising context-dependent gene functions is crucial for understanding the genetic bases of health and disease. To date, inference of gene functions from large-scale genetic perturbation screens is based on ad-hoc analysis pipelines involving unsupervised clustering and functional enrichment. We present Knowledge-Driven Machine Learning (KDML), a framework that systematically predicts multiple functions for a given gene based on the similarity of its perturbation phenotype to those with known function. As proof of concept, we test KDML on three datasets describing phenotypes at the molecular, cellular and population levels, and show that it outperforms traditional analysis pipelines. In particular, KDML identified an abnormal multicellular organisation phenotype associated with the depletion of olfactory receptors and TGFβ and WNT signalling genes in colorectal cancer cells. We validate these predictions in colorectal cancer patients and show that olfactory receptors expression is predictive of worse patient outcome. These results highlight KDML as a systematic framework for discovering novel scale-crossing and clinically relevant gene functions. KDML is highly generalizable and applicable to various large-scale genetic perturbation screens.

## Introduction

Despite the deluge of acquired datasets with High Throughput Gene Perturbation Screening (HT-GPS), the function of a large number of human genes remains poorly understood (Dey *et al*, 2015). Moreover, Gene Ontology (GO), the most comprehensive and structured annotation of gene functions, is largely limited to cell type- and context-independent gene functions(Huntley *et al*, 2015). However, gene function is highly contextual, even for unicellular organisms (Liberali *et al*, 2014; Radivojac *et al*, 2013). Therefore, there is a pressing need for new methods that allow data-driven and context-dependent functional gene discovery based on more complex phenotypes of multi-cellular organisms.

Although HT-GPS has proved to be a powerful method for discovering novel gene functions, the analysis of these datasets has remained a challenging task. This is due to the complexity of phenotypes that the perturbation of a single gene can lead to, as a gene can participate in different functions at different scales. These functions depend on the gene product localisation in the cell (e.g. cytoplasm versus nucleus for transcription factors), cell cycle state (e.g. G1, G2 or S-phase), cell type, cell-cell and cell-microenvironment interactions, and treatment conditions. Existing analysis pipelines based on unsupervised clustering do not generally account for these factors. Consequently, resulting phenotypic clusters are difficult to interpret as they might be composed of different subphenotypes. These challenges are often avoided, particularly in image-based screens, by analysing only a small fraction of the information contained in HT-GPS datasets (Singh *et al*, 2014) which greatly underutilises their potential.

Supervised machine learning and AI systems have been applied successfully in many HT-GPS studies (Held *et al*, 2010; Neumann *et al*, 2010; Shariff *et al*, 2010; Sullivan *et al*, 2018; Eraslan *et al*, 2019). One attractive solution for addressing the lack of phenotypic annotations is the utilization of existing biological knowledge to build intelligent systems that can identify functionally relevant features and phenotypes. Approaches that utilise existing functional annotations have been successfully applied to inference of pathway activity (Schubert *et al*, 2018) as well as prediction a protein function from multiple data types including protein sequence and structure, phylogeny, as well as interaction and gene co-expression networks (Jiang *et al*, 2016; Dey *et al*, 2015; Radivojac *et al*, 2013). Additionally, pioneering work has been done in inferring data-driven gene ontology in yeast (Ma *et al*, 2018; Yu *et al*, 2016; Kramer *et al*, 2014). However, to our knowledge this approach has not been applied in the context of large-scale HT-GPS datasets in multicellular organisms where genetic redundancy and phenotypic complexity are much higher.

Image-based screens are particularly advantageous for inference of biological functions as they provide spatial and context information at single-cell level which allow capturing the emergent behaviours in biological systems (Lock & Strömblad, 2010). Single-cell data are critical for identifying loss-of-function phenotypes that are dependent on cellular state or manifest in only a small sub-population of cells (partial penetrance) (Sacher *et al*, 2008). However, even for a widely-used marker such as DAPI, phenotypic information on nuclear morphology and organisation of cells has not been fully utilised, except in the context of apoptosis and the cell cycle (Neumann *et al*, 2010). The importance of studying the functional relevance of nuclear morphology and multicellular organisation is underlined by the fact that this information is successfully used by pathologists for patient diagnosis based on hematoxylin and eosin stained tumour sections (He *et al*, 2012; Uhler & Shivashankar, 2018). Comprehensive analysis of changes in cell morphology and microenvironment following perturbation is crucial for the identification of genes associated with these important biological traits.

Systematic evaluation of gene sets in biological contexts different from the one in which they are known to function can provide valuable insights into the regulation of biological systems. For example, the roles of genes that regulate developmental processes, such as mesoderm development (MSD), are often not clear in adult tissues. MSD involves the coordination of cell migration, cell adhesion, and cytoskeletal organisation through TGFβ and WNT signalling. The latter pathways are often deregulated in colorectal cancer (Klinowska *et al*, 1994; McMahon *et al*, 2010; Kiecker *et al*, 2016). Another example is the olfactory receptor family, which were discovered in 1991 in sensory neurons and are composed of about 400 genes in humans. Olfactory receptors have been shown to be expressed in many non-sensory tissues in both development and adult tissues including the gastrointestinal system (Maßberg & Hatt, 2018). However, we have little understanding of their functions in these contexts.

Here we propose KDML, a novel framework for automated knowledge discovery from large-scale HT-GPS. KDML is designed to account for pleiotropic and partially penetrant phenotypic effects of gene loss. We apply this framework to three large-scale datasets generated by different methods, describing phenotypes at the molecular, cellular and tissue levels and show it outperforms existing analysis pipelines. We analyse a cell organisation phenotype that KDML identifies and links to genes annotated to the Mesoderm Development (MSD) term. KDML predictions include many genes in TGFβ and WNT signalling pathways as well as many olfactory receptors. Through an integrative analysis with gene expression data of colorectal cancer patients, we validate the link between expression of olfactory receptors and TGFβ and WNT signalling, and show that the expression of some olfactory receptors can stratify the outcome of higher-grade colorectal cancer patients. In summary, KDML is a flexible and systematic framework for comprehensively analysing HT-GPS and identifying context and tissue-dependent gene functions.

## Results

### KDML provides a systematic framework for inferring gene functions from high dimensional phenotypic data

KDML utilises existing biological knowledge to automatically identify gene perturbation phenotypes in HT-GPS datasets and map these phenotypes to potential biological functions. Using GO annotations to define functional annotations, KDML trains independent binary Support Vector Machine (SVM) classifiers for each GO term (Methods). Each classifier searches whether genes annotated to a GO term have a distinct perturbation phenotype in a given dataset (Fig. 1A). Once trained, each GO term classifier can be applied to rank all perturbed genes in the dataset based on their phenotypic similarity to the term’s annotated genes. For example, a classifier of cell cycle function is expected to identify a difference in proportions of different cell cycle states and/or nuclear morphology in gene perturbations annotated to this function. Once a quantitative signature is established for annotated genes, then gene perturbations that have similar signatures will be predicted to participate in the cell cycle. We included terms with a sufficient number of positive examples from the three GO Ontologies: biological process, molecular function, and cellular component (Fig. 1B, Methods, and Supplementary Table 1). On average, each gene has 30 annotations (Fig. 1C) with 2,152 genes having more than 50 annotations and 2,908 genes having no annotations based on the selected terms.

**Figure 1.**
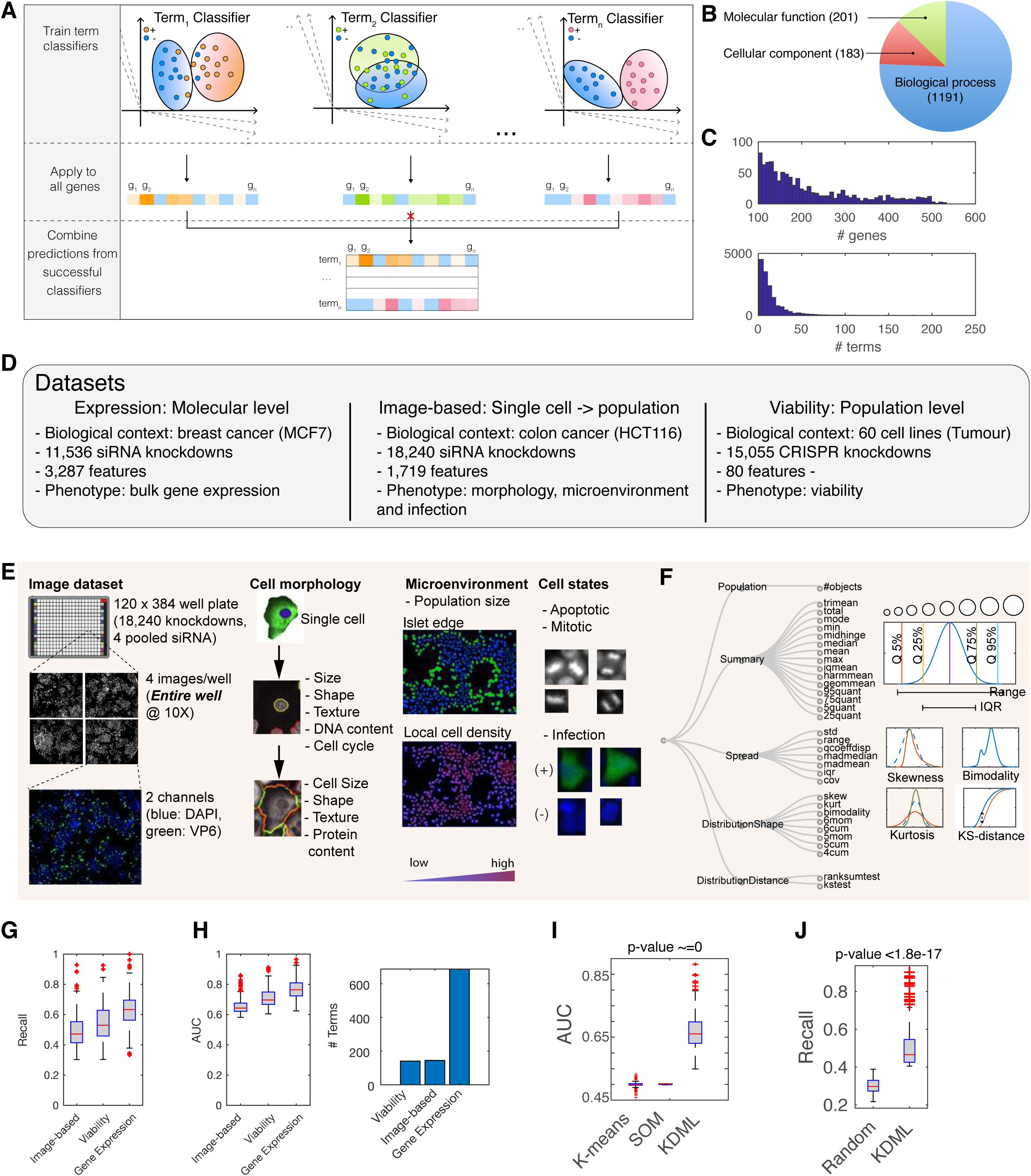
Training and predicting gene functions from HT-GPS. (A) KDML workflow. Feat: feature. (B) Tested datasets. (C) Overview of single cell-resolved image-based features measuring morphology, microenvironment and infection (C) and the generated statistics based on a population of single-cells when applicable (D) (n>625 and Supplementary Table 2). Examples on the various measurement types are shown on the right and include various quantiles, mean, range, IQR (interquartile range), kurtosis, skewness, bi-modality and KS distance. (E) Categories and numbers of included GO terms. (F) The distribution of genes per term and vice versa based on GO annotations. (G) AUC and recall of test samples of classifiable terms across the three tested datasets. (H) The number of classifiable terms in the three datasets. (I) The number of shared genes for the most overlapping terms across datasets. (J) Selected features by ncRNA processing classifier and their average values in predicted genes based on the viability dataset. Viability for colon and breast cancer cell lines is indicated in red and green respectively. (K-L) Benchmarking of KDML against (K) clustering of genes using k-means or SOM (self-organising maps) or (L) classifiers that were trained to classify random sets of 200 genes. Box plots elements: centre line, median; box limits, 25^th^ and 75^th^ percentiles; whiskers, +/−2.7 standard deviation. Points: outliers.

We tested KDML on three HT-GPS datasets: 1) pooled genome-wide CRISPR/Cas9 screens measuring cell viability in 60 cancer cell lines(Rauscher *et al*, 2018), 2) a large-scale siRNA screen measuring changes in the expression of 3,287 genes in MCF7 breast cancer cells(Duan *et al*, 2014), and 3) an arrayed image-based genome-wide siRNA screen measuring changes in 168 single-cell features that quantify morphology, microenvironment, and infection in HCT116 colorectal cancer cells based on stains of DAPI and the rotavirus-expressed Viral Protein 6 (VP6) (Green & Pelkmans, 2016) (Fig. 1D-E). The latter dataset is composed of single-cell measurements, in which each perturbed population has 6,040 cells on average (Supplementary Fig. 1A). To capture the heterogeneity in cellular states and collective cellular behaviour as readouts (Altschuler & Wu, 2010), we aggregated single-cell features into 1,719 features per gene perturbation profile. Computed statistical measures include standard deviation (SD), various quantiles, skewness, kurtosis, bimodality coefficient, Kolmogorov Smirnov (KS) and rank sum statistics (Fig. 1F, Methods and Supplementary Table 2). Furthermore, we corrected all features for their dependence on cell number which reduced classifier bias toward gene perturbations with a very low or high number of cells (Supplementary Fig. 1A-D and Methods).

For each of the datasets, we selected the most confident GO term classifiers based on the recall (true positive rate) and false positive rate such that it is significantly better than classifying a random set of genes (Methods). The HT-GPS dataset using gene expression as a readout had the highest recall and Area Under the Curve (AUC, which reflects specificity versus sensitivity) as well as the number of classifiable terms (Fig. 1G-H). This is expected since this dataset measures for each perturbation changes in the expression of 3,287 genes, whose variation is representative for most of the transcriptome (Duan *et al*, 2014). Interestingly, viability measured in 60 cancer cell lines in different contexts is also highly informative on gene functions with a comparable number of classifiable terms to the screen based on multivariate image-based readouts (Fig. 1G-H).

Multivariate single cell-resolved image-based readouts outperformed gene expression for some GO terms while 33 terms were only classifiable based on image-based single-cell features (Supplementary Fig. 1F-G). Many of those are involved in phosphorylation, ubiquitination or membrane transport. This might be due to the fact that bulk gene expression does not necessarily capture post-transcriptional events or measure changes at the single-cell level. For example, exopeptidase activity and voltage-gated channels had a higher overall recall based on image-based single-cell features compared to the gene expression dataset (Supplementary Fig. 1E). These results show that different experimental techniques for probing biological systems complement each other and provide different functional information.

We compared KDML against commonly used analysis pipelines based on dimensionality reduction and unsupervised clustering. We clustered gene profiles using k-means and self-organizing maps while varying the number of clusters (Methods). KDML significantly outperformed clustering approaches, with the latter scoring close to a random AUC of 50% (Fig. 1I). Additionally, KDML performed significantly better than random sets of genes (Fig. 1J, p-value <1.8443e-17 and Methods).

Below, we focus on the further analysis of the image-based perturbation dataset, as it is the only dataset that provides spatially resolved single-cell measurements which are more challenging to analyse and interpret (Collins, 2009).

### Using KDML to identify GO terms represented in the image-based dataset and their relations

To illustrate the use of KDML, we apply it to identify the functional information in the image-based single-cell perturbation dataset. We trained KDML using a subset of features to determine the GO terms that can be learned from different cellular markers (morphology and microenvironment features based on DAPI versus infection features based on VP6 staining) (Methods). Shape features are predictive of 61 GO terms while infection measurements are predictive of 39 terms, with 8 terms shared (Fig. 2A and Supplementary Fig. 2A). Combining both feature sets, result in an additional 54 classifiable terms (Fig. 2A). These results demonstrate that multivariate imaging data can be predictive of many gene functions even when only one or two cellular stains are used.

**Figure 2.**
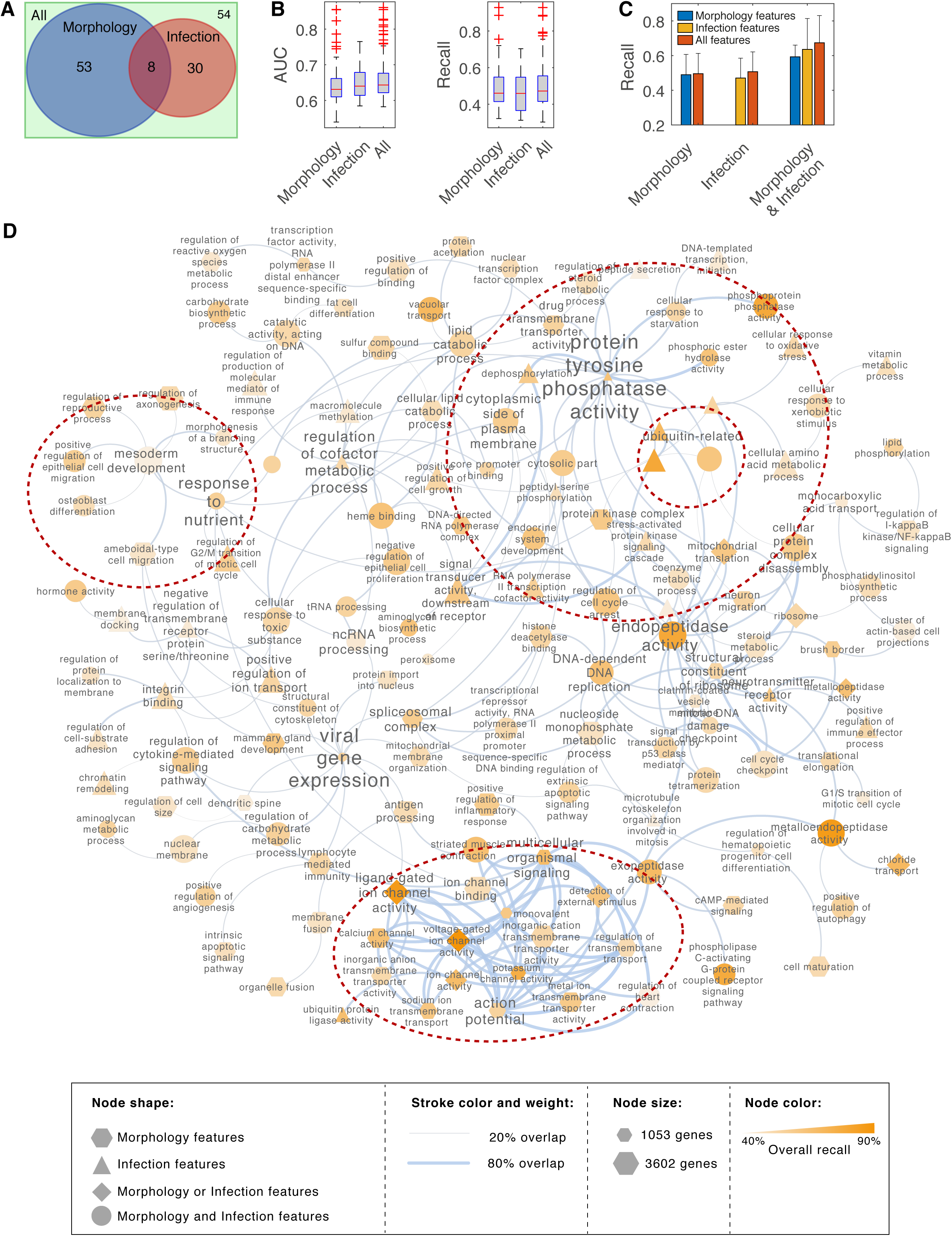
Analysis of functional information using KDML based on image-based dataset. (A) The number of classifiable terms when using morphology versus infection features. (B) AUC and recall of test samples for classifiable terms when using different subsets of features. Box plots elements: centre line, median; box limits, 25^th^ and 75^th^ percentiles; whiskers, +/−2.7 standard deviation. Points: outliers. (C) The improvement in performance when all the features are used for terms that are classifiable when only shape features are used, only infection features are used, or either shape or infection features are used. Only a slight improvement in classification performance is achieved by combining feature subsets when the signatures based on a subset of features are already strong (D) Network representation of classifiable terms where edges indicate the overlap in the predicted gene lists between GO terms.

To gain insight into the functions that have been learned by KDML, we generated a network of classifiable GO terms based on the overlap in their predicted gene lists (Fig. 2D). We observe a strong cluster of membrane transport-related terms including potassium, calcium, sodium and metal ion channels (Fig. 2D). Most of these terms can be classified using morphology and microenvironment features alone and their phenotypic profiles cluster together (Fig 3A, C1 and Supplementary Fig. 3).

**Figure 3.**
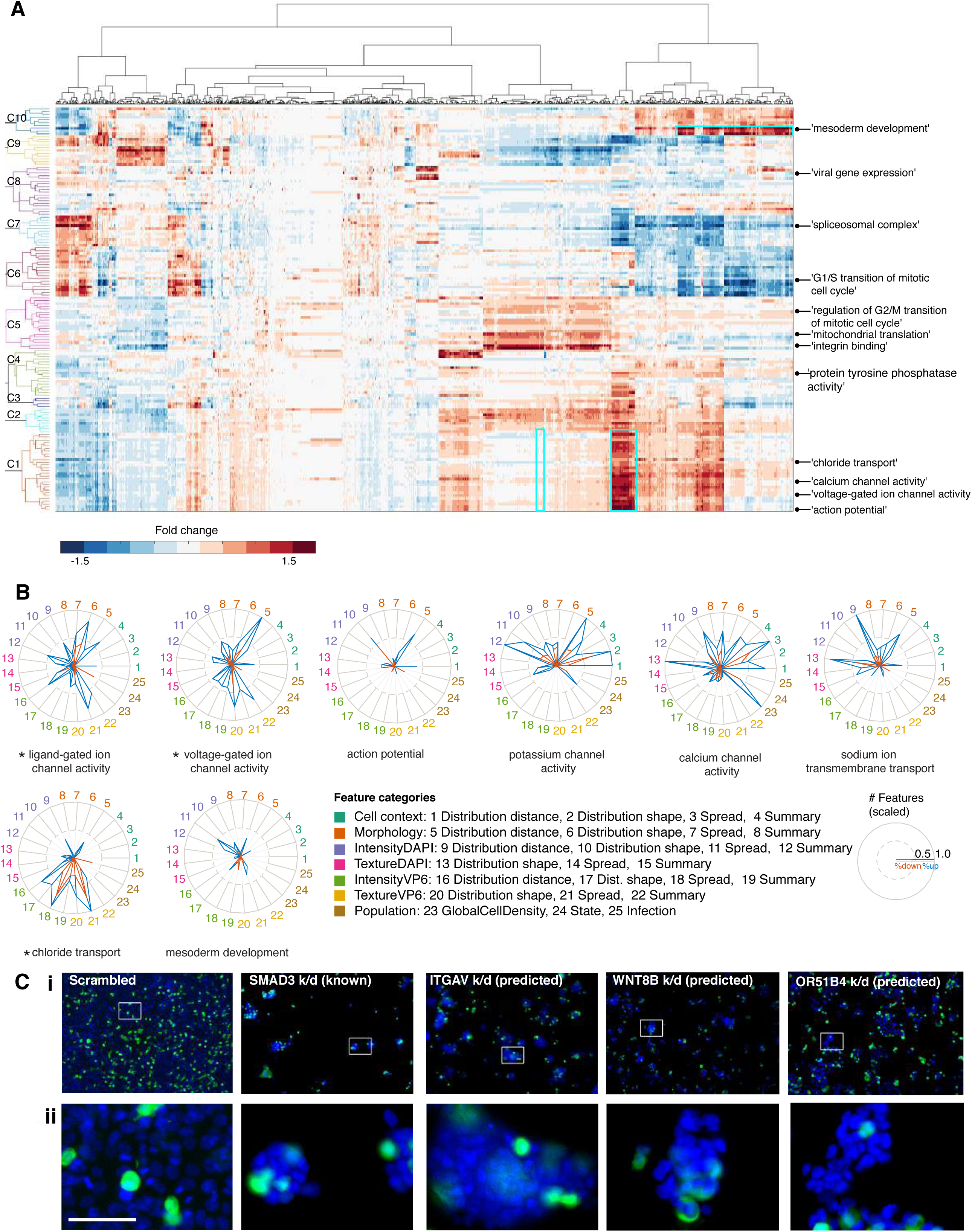
Association between phenotypic changes in single cell-based features and GO terms. (A) Hierarchical clustering of the average fold change of predicted positive versus negative samples for each term using all features. Hierarchical clustering is based on ward linkage and Euclidean distance. Regions outlined by cyan rectangles are shown in Supplementary Fig. 3-4A. (B) The number of features in different categories that are selected by the respective GO term (scaled). Blue indicates the number of features with a higher average than control while red indicates the number of features with a lower average than control. * indicates the GO terms that are classifiable by either shape or infection features. (C) Example cell images following knockdown (k/d) of known or predicted MSD genes versus control (scrambled). ii) zoom-in image of the region highlighted in (i). Blue: DAPI, and Green: VP6 antibody. Scale bars = 65 µm.

Interestingly based on our data, ion channel terms (molecular level) are linked to multicellular organismal signaling function (tissue level). Another cluster included many phosphorylation- and ubiquitination-related terms, many of which are classifiable based on infection features (Fig. 2D). Therefore, KDML is able to classify biological functions spanning from the molecular-to the tissue-level.

### KDML automatically maps GO terms to functional phenotypes

To understand the phenotypic changes associated with different GO terms, we categorised the phenotypic features based on feature type (cell context, morphology, DAPI intensity and texture, VP6 intensity and texture, state, etc.) as well measurement type (summary statistics, spread statistics, distribution shape or distribution distance - the distance between perturbed and control distributions) (Supplementary Fig. 2E and Supplementary Table 2). Next, we counted and scaled the number of features selected by a term classifier in the different categories (Fig. 3B).

Membrane transport-related terms result in an increased number of cells and total cell area as well as local cell density and cell distance to islet edge (Supplementary Fig. 3B). Interestingly, the ‘ligand gated ion channel activity’ and ‘voltage gated ion channel activity’ terms are the most accurately classified (Supplementary Fig. 2A). These terms are classifiable based on either shape features or infection features but Infection features were almost dispensable for their classification when all features were considered (Supplementary Fig. 2D). Predictive features of these two terms include distribution shape and distribution distance of cell morphology (Fig. 3B, FeatCat 5-6). Classifiers of calcium and potassium channel activity selected many more microenvironment features than gated channels (FeatCat 1-4). Notably, the rank-sum statistic was among the most selected measure by different classifiers indicating its biological relevance and robustness (Supplementary Fig. 2E). These results illustrate the functional relevance of our distribution measures, and also that single-cell aggregated measurements of cell morphology and context can be highly predictive of ion channel-regulating genes.

Although chloride transport is also classifiable by morphology or infection features, shape-features are dispensable for its classification (Fig. 3B and Supplementary Fig. 2D). Depletion of chloride transport genes affected many texture and intensity measures of VP6 but most interestingly it significantly decreased rotavirus infection (p-value <4.9e^−139^, Fig. 3B and Supplementary Fig. 3A,C). The association between chloride transport genes and rotavirus is consistent with the fact that rotavirus increases chloride secretion (Lorrot *et al*, 2006; Yin *et al*, 2017). Our results now suggest that chloride channels might be required for the spread or replication of rotavirus, implying that increased chloride secretion is not just a downstream consequence of virus infection, but a means of the virus to spread.

MSD genes are predicted based on the spread and summary values of cell context, DAPI intensities, as well as the distribution shape of cell morphology, suggesting changes in cell organisation (Fig. 3B and Supplementary Fig 4A). Indeed, depletion of previously annotated (e.g. *SMAD3*) and predicted (e.g. *ITGAV*, *WNT8B* and *OR51B4*) MSD genes results in small cell colonies compared to control (Fig. 3C). This phenotype might indicate the inability of cells to spread or migrate. This hypothesis is supported by the high overlap of the MSD predicted genes with GO terms such as migration (epithelial and ameboidal-like), morphogenesis (e.g. morphogenesis of branching structure and axonogenesis), and chemotaxis (response to nutrients) (Fig. 2D). Thus, our holistic analysis reveals a role for MSD-associated genes in the spread and organisation of epithelial colorectal cancer cells into a uniform epithelial sheet.

### KDML allows pleiotropic analysis of high dimensional data

KDML allows pleiotropic analysis by training an independent classifier for each GO term where each classifier can be considered as an expert in a given biological function. These classifiers can rank all genes based on phenotypic similarity. For example, previously annotated genes to MSD are predicted to perform various additional functions such as positive regulation of epithelial cell migration, and mammary gland development (Fig. 4A). Interestingly, MSD genes that are predicted to play different sub-functions tend to occupy different regions in the MSD sub-phenotypic space which is reflected by differences in a subset of MSD features (Fig. 4B and Supplementary Fig. 4B). The pleiotropic nature of gene function allows subspecialisation as genes perform a different combination of functions.

**Figure 4.**
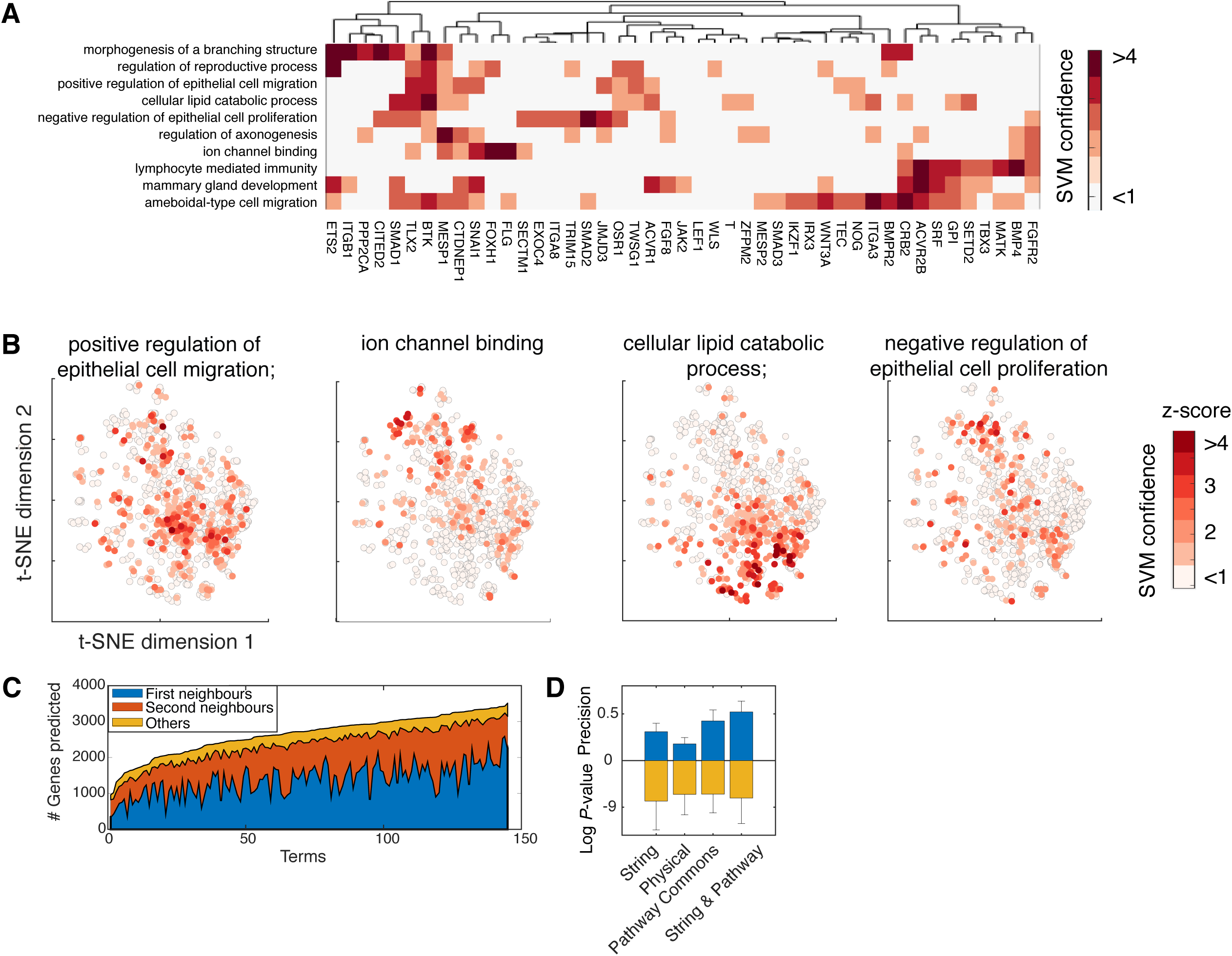
KDML allows pleiotropic analysis of high dimensional phenotypic data. (A) Heatmap showing SVM-based ranks (z-scored) for known MSD genes against multiple functions. The functions on the y-axis are the top ten overlapping terms with MSD classifier prediction. (B) Embedding of MSD subphenotypic space using t-SNE based on the selected features by MSD classifier where only MSD genes are considered (Methods). Colour indicates the respective SVM rank for each gene for the corresponding function. (C) The number of genes that are positive for a given GO term and the extent of their interactions with the annotated genes for that term either as first or second neighbours. Interactions are based on STRING and Pathway Commons databases. (D) Predictions of many GO terms are significantly enriched for protein-protein interactions based on the number of first-neighbour interactions based on STRING, physical (based on experimental evidence in STRING), and Pathway Commons (right tail Fisher Exact Test). Error bars indicate 1 standard deviation.

Perturbation of a biological function might affect only a subset of the measured features. For example, the mean and integrated intensity of DAPI is significantly lower in genes predicted to participate in cell cycle checkpoint (Supplementary Fig. 5A). However, these genes do not cluster together when all phenotypic features are considered, but rather only when considering features selected by cell cycle checkpoint classifier (Supplementary Fig. 5B-C). Identification of sub-phenotypic effects associated with each gene allows determination of which of those sub-phenotypic effects are potential off-target effects. Predictions of genes that are targeted by the same siRNA seed (guide strand) are filtered out if the seed is significantly over-represented in a given GO term classifier (Supplementary Fig. 5D and Methods). This further underlines the importance of searching in the sub-phenotypic space to deconvolute biological signals contained in high dimensional data and obviate off-target effects.

Pleiotropy and functional subspecialisation are achieved via protein-protein interactions. We found that most of predicted genes for a GO term are interacting based on String and Pathway Commons (Fig. 4C). Many of these genes are either first or second neighbours in the interaction network of genes previously annotated to that term (Fig. 4C-D, and Methods). This provides further validation of KDML predictions as it is expected that the depletion of interacting genes results in a similar phenotype.

### Validation of mesoderm development classifier predictions in the context of colorectal cancer

Since the loss of tissue organisation is one of the main characteristics in tumours (Hinck & Näthke, 2014), we sought to analyse MSD genes whose perturbation resulted in abnormal cell organisation in HCT116 cells. MSD genes are significantly enriched for morphology-regulating pathways including tight junctions, focal adhesion, and cell cytoskeleton (Fig. 5 and 6A and Supplementary Table 4). This is expected, as cell adhesion and actin cytoskeleton are required for cell spreading and motility. Among the predicted genes are 15 collagen, 12 integrin and many polarity genes, such as Par3 and Par6 (Supplementary Table 4). Surprisingly, many of the predicted MSD genes also participate in pathways that are often deregulated in colorectal cancer including TGFβ, WNT and PI3K-AKT pathways (Muzny *et al*, 2012) (Fig. 5). Predicted genes include *TGFβR2*, *PTEN* and *ERBB2* which are often mutated in cancer (Kuipers *et al*, 2015). Over-activation of TGFβ and WNT is associated with mesenchymal and stemness phenotypes in colorectal cancer respectively (Hinck & Näthke, 2014). But our results suggest that genes in these pathways are also required for the formation of monolayer epithelial sheets.

**Figure 5.**
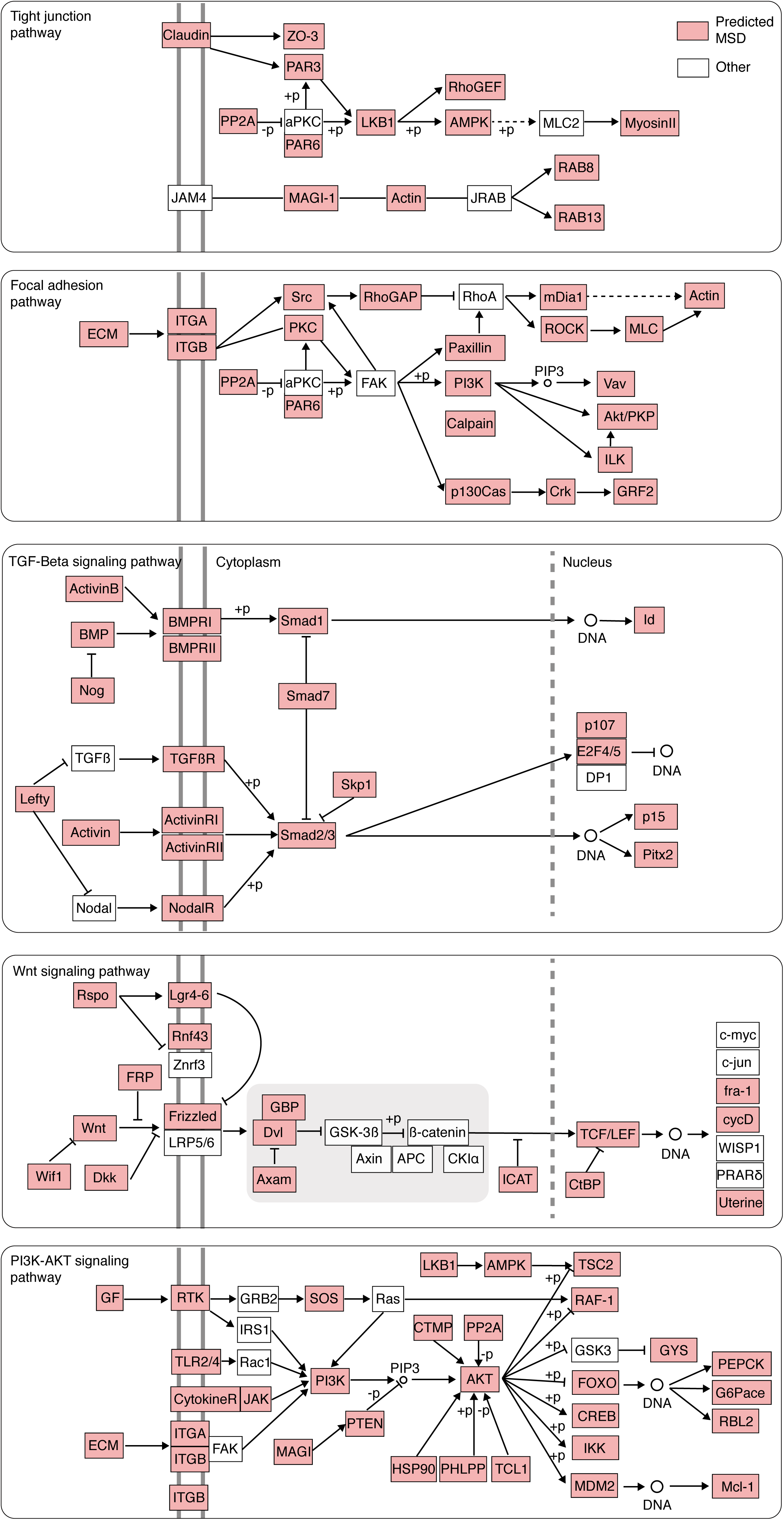
Predictions of MSD classifier are significantly enriched for cell adhesion and colorectal cancer pathways. Snapshots from tight junctions, focal adhesion, TGFβ, WNT, and PI3K-AKT KEGG pathways where MSD genes are highly represented.

**Figure 6.**
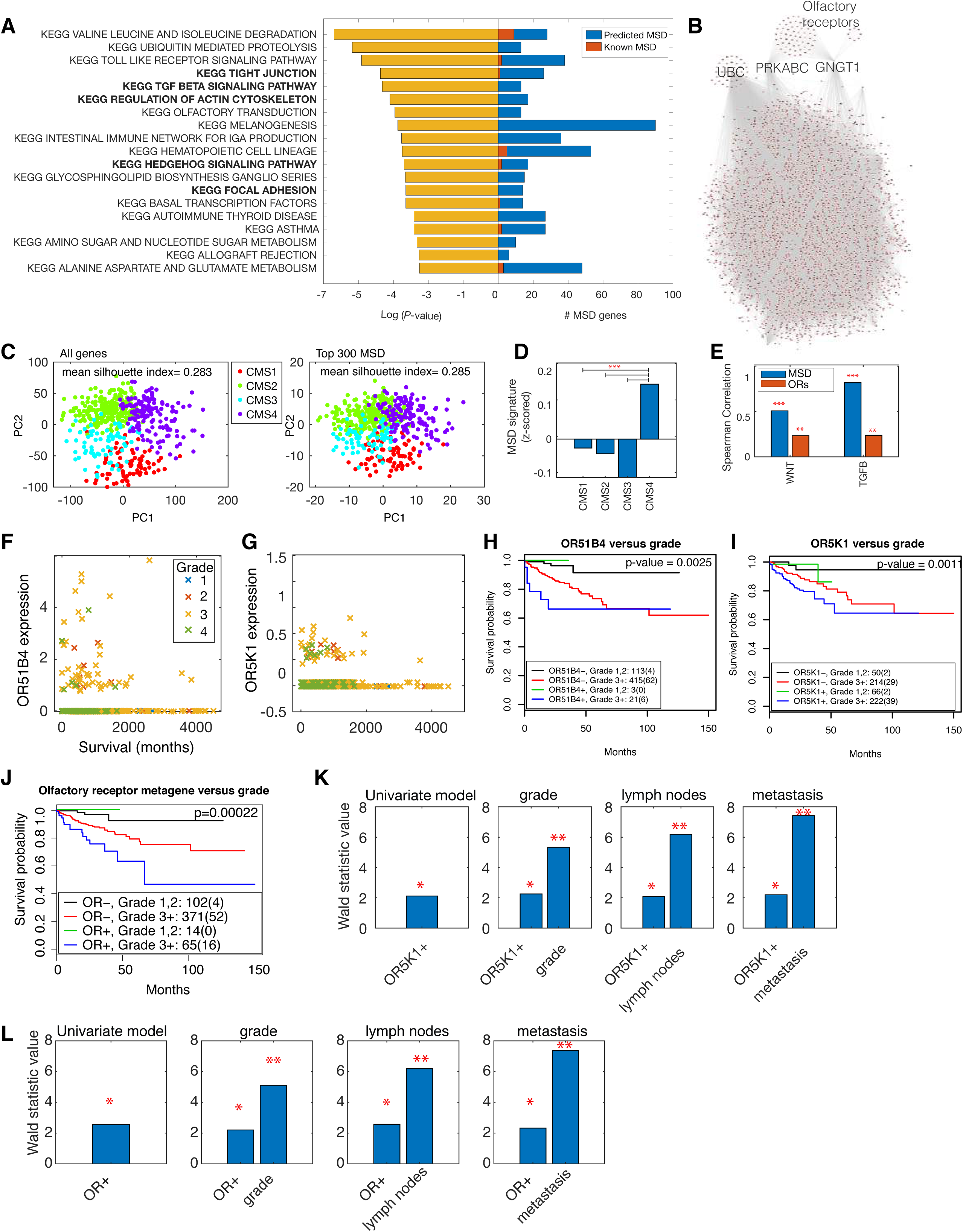
Analysis of MSD predicated gene list. (A) KEGG pathways that are significantly represented in MSD genes (p-value<0.05 based on right tail Fisher Exact Test). Previously annotated genes (known) for MSD are shown in red while others are predicted by KDML based on phenotypic similarity to known MSD genes. (B) Network depicting the interactions between MSD genes based on STRING and Pathway Commons. (C) TCGA colorectal cancer patients’ data projected into the first two principal components based on the expression of all genes (left) and expression of the top 300 predicted MSD genes (right). (D) Average expression of the top 300 predicted MSD genes across colorectal cancer molecular subtypes (CMS1-CMS4) (*** p-value < 3.7×10^−9^). (E) Spearman correlation coefficient between average top 300 MSD genes or average MSD-associated olfactory receptors and TGFβ or WNT signatures (p-value<0.0001 and < 0.05 respectively). ORs: olfactory receptors. (F-G) Survival of colorectal cancer patients (months) against the expression of *OR51B4* (F) and *OR5K1* (G). Colour indicates tumour grade where grade 1 is well differentiated, grade 2 is moderately differentiated and grade 3-4 is poorly differentiated. (H-J) Kaplan Meier survival analysis of colorectal cancer patients based on Grade 3+/− tumours and the expression state of *OR51B4* (H), *OR5K1* (I) and olfactory receptor metagene (J), which is aggregated, based on the expression of many olfactory receptors (Methods and Supplementary Fig. 6E). (K) Wald statistic value based on Cox Proportional Hazard regression analysis of *OR5K1* predictivity of survival against tumour grade, presence of lymph nodes and metastasis. Detailed results for all tested variables and model significance are shown in Supplementary Table 6. * indicates p-value<0.05 and ** indicates p-value <0.001.

Collectively, this suggests that predicted MSD genes might play a role in colorectal cancer by regulating cell organisation via TGFβ and WNT signalling.

To validate the association between predicted MSD genes and colorectal cancer, we interrogated The Cancer Genome Atlas (TCGA) gene expression dataset (Muzny *et al*, 2012) of 575 colorectal cancer patients. First, we sought to investigate whether MSD genes can reflect the consensus molecular subtypes (CMS) of colorectal cancer: CMS1 (microsatellite instability), CMS2 (WNT activation), CMS3 (metabolic), and CMS4 (mesenchymal) (Guinney *et al*, 2015). Strikingly the top 300 predicted MSD genes recapitulate colorectal cancer molecular subtypes with comparable performance to using all genes (Fig. 6C, Methods, and Supplementary Table 5). This is significantly higher than random sets of 300 genes (p-value = 3.4389e-25 and Methods). Furthermore, the average expression value of the top 300 MSD genes is significantly higher in CMS4 subtype (Fig. 6D), which is associated with TGFβ signalling and predictive of worse prognosis.

Next, we sought to validate the relationship between TGFβ and WNT signalling, and MSD genes in colorectal cancer. Average expression of WNT genes is used as a surrogate for WNT signalling while TGFβ signature is based on genes reported by a previous study (Calon *et al*, 2012) (Methods). We found a significant correlation between MSD genes and TGFβ and WNT signatures in colorectal cancer patients (Fig. 6E). These results suggest coordination between MSD genes and TGFβ/WNT signalling during tumorigenesis.

Strikingly 84 olfactory receptors are predicted by MSD classifier (Fig. 6B and Supplementary Table 6). These also significantly correlate with WNT and TGFβ signatures (Fig. 6E). We sought to investigate the correlation between olfactory receptors and colon cancer patient outcomes as they had only a few connections to the rest of MSD network making them potential therapeutic targets (Fig. 6B and Supplementary Fig. 6A). We generated a metagene representing the state of many of olfactory receptors as they are expressed in a small number of patients (Methods and Supplementary Fig. 6B-E). Expression of many olfactory receptors, albeit at a low level, correlates with higher tumour grade and predictive of significantly worse patient outcome (Fig. 6F-J and Supplementary Fig. 6E-H). These include *OR51B4* and *OR5K1* that are overexpressed in HCT116 cells (Barretina *et al*, 2012). Moreover, expression of *OR5K1* or olfactory receptor metagene is predictive of survival, independent of other clinical variables including tumour grade, presence of lymph nodes, and metastasis state (Fig. 6K-L and Supplementary Table 7-9). Altogether, KDML can identify clinically relevant and disease-specific genes as it systematically links gene perturbation phenotypes to functions while minimising human biases in the analysis pipelines.

## Discussion

Intelligent machine learning methods that integrate existing biological knowledge in a systematic fashion are crucial for accommodating the explosive growth of phenotypic data at multiple system levels. Here we propose a computable and flexible framework that integrates functional annotations to automatically identify three-way relationships between phenotypes, genes, and biological functions. Not only does KDML outperform clustering-based approaches, but it also accounts for the pleiotropic nature of gene function and can mitigate the problem of off-target effects. KDML can be applied to various phenotypic readouts including image-based and gene expression datasets.

A general problem in inferring functions from HT-GPS is that the gene perturbation phenotype might be a result of affecting a biological function directly or indirectly through participating in a related function. Perturbation effects, can then propagate through the protein-protein interaction networks. Indeed, we observe a high enrichment of protein-protein interactions for genes that are predicted to share a given term. This may explain the high number of positive predictions, as many neighbours in the interaction network tend to have similar subphenotypes. Predictions can be prioritised based on SVM confidence or against other biological annotations such as protein-protein interactions or KEGG pathways. Alternatively, high confidence predictions can be obtained by integrating results from various datasets. Taken together, our predictive analysis framework serves as a tool to generate testable hypotheses and pave the way to more integrative studies.

One limitation of our approach is the dependency on noisy GO annotations which do not provide perfect ground truth. Moreover, a sufficient number of positive genes is required to classify a certain biological function. Both of these factors can restrict the application of data-hungry deep learning methods and might influence KDML performance. Nonetheless, these issues also apply to existing pipelines and will be reduced as our databases of gene function expand.

Cellular morphology has been illustrated to reflect multiple aspects of cell physiology (Fuchs *et al*, 2010). Here we further show that advance measures of single-cell distributions in image-based screens are useful for identifying genes regulating multicellular functions. Notable, only cellular populations with at least 675 cells were considered in our analysis which might not be feasible to achieve using single cell sequencing technologies. Importantly, phenotypes based on collective cellular behaviour can be difficult to discern by human eye. KDML identified that depletion of annotated ligand- or voltage-gated channel genes affects cell area and microenvironment measures at the population level. This is consistent with previous reports on the role of ion channels in cell volume regulation and proliferation (Lang *et al*, 2007). At a higher system level, KDML predictions for ion channel terms overlapped with the multicellular organismal signalling. This implicates that ion channel perturbations affected cell-cell communication akin to their function in neurons (Barshack *et al*, 2008; House *et al*, 2015) but further experiments are required to validate that. Importantly, aggregated measures of single cell features outperformed gene expression data in the identification of many membrane-transport functions.

Our systematic analysis revealed a role of MSD genes in multicellular organisation which manifested in cell clumping and increased local cell density in colorectal cancer cells. Cell microenvironment has been shown to contribute to cancer initiation and progression (Friedl & Alexander, 2011). Our analyses demonstrate that perturbation of many extracellular matrix and integrin genes can result in a similar phenotype to perturbing TGFβ and WNT signalling. This suggests an important link between cell microenvironment (adhesion) and shape (cytoskeleton) and determination of cell fate (i.e. stemness and differentiation via TGFβ and WNT signalling). Consistent with this, SMAD3, which plays an essential role in TGFβ signalling, has been shown to link shape information to transcription in breast cancer cells (Sailem & Bakal, 2017), while WNT signalling can be linked to cell microenvironment via the differential localisation of its downstream effector β-catenin. Therefore, KDML can provide insights on the emergence of subphenotypes, such as cell organisation, based on the integration of different signalling pathways.

The predicted role of olfactory receptors in MSD is consistent with previous reports on their expression in developing mesoderm tissue and their role in patterning (Nef & Nef, 2002; Weber *et al*, 2002; Dreyer, 2002). Moreover, they are associated with non-canonical WNT signalling in neuronal cells (Zaghetto *et al*, 2007) and TGFβ in progenitor neuronal cells (Getchell *et al*, 2002). We show that the expression of *OR51B4* and *OR5K1*, even at a low level, correlate with patient grade and worse prognosis. This supports our results as higher-grade tumours are characterised by poorly differentiated cells and loss of epithelial organisation (Guinney *et al*, 2015). Olfactory receptors in the gut might be activated by odours and chemicals produced by microbes or ingested food. Based on our results this might, in turn, activate dedifferentiation via crosstalk with TGFβ and WNT pathways. We propose that blockade of olfactory receptors might provide a novel therapeutic strategy in colorectal cancer.

In summary, our results illustrate the generalizability and utility of KDML as a framework for systematic gene function discovery from HT-GPS across multiple biological scales. We believe that KDML can scale-up to more complex phenotypic screens utilising advance multiplexing technologies (Gut *et al*, 2018) or probing microtissues or organisms. We envision that systematic application of KDML to the large amounts of generated HT-GPS datasets will greatly accelerate and advance our understanding of gene functions at the molecular, cellular and tissue levels which can lead to the discovery of new therapeutic target genes.

## Methods

### KDML implementation and data analysis

KDML pipeline and all analyses were performed using MatLab (http://www.mathworks.com/) unless stated otherwise.

### Preparation of GO annotations

GO annotations were downloaded from UniProt (Feb, 2018). Annotations of child terms were escalated up to parent terms in order to have a sufficient number of positive examples for classification. Only GO terms that had 100-500 gene annotations were considered resulting in 1575 terms. Highly redundant GO terms (jaccard index>70%) were merged.

### Gene profiles computation for image-based readouts

Image analysis was performed using CellProfiler as previously described(Green & Pelkmans, 2016). Briefly, all images were pre-processed using illumination correction and background subtraction. Nuclei were segmented based on DAPI channel. Cells were defined as 10-pixel expansion of the nucleus. Shape, intensity and texture features were quantified for DAPI and VP6 channels. SVM classifiers were trained to identify mitotic, apoptotic, poorly segmented and infected cells. Infection index was corrected for population context(Green & Pelkmans, 2016). The number of cells in different states was normalised to the number of cells in the well.

Single-cell data were aggregated per well using various statistical measures (Supplementary Table 2). KS and rank sum statistics were used to compare a treated population to scrambled/empty populations (distributions distance).

The rank-sum statistic was computed based on Mann-Whitney-Wilcoxon rank sum test using ranksum MatLab function. All values of a feature X were ranked when combining perturbed and control samples. The sum of the ranks of the perturbed sample was then used to calculate the statistic as following

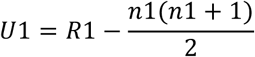

where U1 is the rank sum statistic, R1 is the sum of cell ranks in the perturbed sample of cells and n1 is the size of the perturbed sample.

KS distance was computed as the KS statistic based on a two-sample Kolmogorov-Smirnov test using kstest2 MatLab function. KS distance measures the maximum distance between the cumulative distribution of perturbed and control cell populations.

Both KS statistic and ranksumtest statistic were multiplied by a scaling factor *sf* to account for differences in sample sizes.

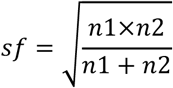

where n1 is the size of the perturbed sample and n2 is the size of the control sample.

### Feature subsets of Image-based dataset

All features based on DAPI channel as well as cell context measures were defined as morphology features. On the other hand, features based on VP6 channel including texture and intensity were defined as infection features. These also include the microenvironment of the infected cells such as how many infected cells on the islet edge.

### Data pre-processing

All datasets were cleaned by excluding samples or features with more than 30% missing data. Remaining missing values were imputed based on the weighted mean of the nearest 10 neighbouring features based on Euclidean distance.

- siRNA gene expression: siRNA gene expression profiles were averaged per gene. Principal component analysis was performed and the first 1000 principal components (z-scored) were used for KDML classification.
- Viability: Gene perturbations or cell lines where more than 30% of their values were missing were filtered out. Then viability scores were z-scored.
- Image-based datasets: Genes that significantly reduced viability (<625 cells) are filtered to avoid SVM bias toward viability phenotypes(Green & Pelkmans, 2016). All data were z-scored by subtracting the plate mean and dividing by the plate Standard Deviation (SD). Then the average per gene perturbation was computed. To eliminate features dependency on cell number, cell number was binned into 32 bins such that each bin had at least 100 gene perturbation profiles. Finally, feature values for samples in each bin were z-scored to the corresponding bin mean and SD.

### KDML training

Training of classification models: The same training pipeline was applied to the three datasets. The annotated genes for each GO terms were split into 70% for training and 30% for testing. Different classification methods were tested on a subset of terms including Random Forests, Lasso Regression and SVM. SVM was chosen because it resulted in less overfitting. A binary SVM with a Radial Basis Function (RBF) kernel was trained per term to classify the annotated genes (positive class) against a set of randomly selected genes excluding annotated genes (negative class). SVM is trained based on a balanced number of genes in the negative and positive class. All training was performed with a 30 fold cross validation for training samples to avoid over-fitting. Scale of SVM kernel (sigma) is optimised by brute force search based on F-score metric. F-score consider sensitivity as well as specificity of the model by computing the harmonic mean of precision and recall metrics as following:

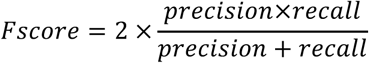

Feature selection: Forward feature selection was applied to identify discriminative features for each term. Features were initially sorted based on KS test p-value comparing positive and negative samples. Sorted features were added sequentially to the model and only features that improved the model performance based on F-Score were retained. No more than 100 features were allowed per model.

Restricting KDML to gene annotations based on experimental evidence codes in UniProt (EXP, IDA, IEP, IMP, IGI, and IPI) was tested but did not improve the performance. We also tested excluding genes in terms that are semantically similar to the term under classification from the negative class. This also did not have a significant effect on KDML performance as selecting these genes had a very low probability when a random sample of all genes was selected.

### Selection of classifiable GO terms

The classification performance can be affected by multiple factors: 1) The quality of GO annotations. 2) For a given GO term, not all annotated genes would perform that function in the investigated cell type or context. 3) Incomplete data: the set of phenotypic readouts acquired by a certain experiment provide a partial description of cellular states. 4) Perturbations efficiency.

Threshold estimation for selecting classifiable GO term: the cut-off for recall on test samples was set to 30% as all classifiers of random gene sets scored less than this cut-off when no more 20% of genes were predicted positive and at least 40% of training samples are classified correctly.

Random gene sets: 100 random gene sets were drawn with each sample having 200 genes. Then KDML was trained to classify these random samples.

### Comparison between KDML and unsupervised analysis methods

Principal component analysis was applied to reduce the dimensionality of the data. K-means and self-organising maps clustering methods were applied to the first 100 principal components which captured 96.25% of the variability in the data. The number of clusters was varied between 10 and 200 clusters, as it is not trivial to determine the number of clusters. Hierarchical clustering was also tested but found to result in one big cluster and many clusters with very few genes.

### Generation of GO term network for Image-based dataset

The classifiable GO terms based on the image-based dataset are connected by an edge if the overlap in their predicted gene lists is greater than 70% based on Jaccard Index. GO terms that do not score an overlap above this threshold are connected to the term with the highest overlap. This is performed iteratively until all the terms are connected to the network. The generated network is visualised in Cytoscape v3.3.

### Interaction enrichment

For a given GO term classifier, we counted the number of positive and negative genes on one hand, and those that are first neighbours of the annotated genes for that term or not. Then, Fisher exact test was used to calculate the significance of the number of first neighbour interactions.

### Detection of seed effects

A pool of four siRNA was used in the image-based single cell resolved dataset. The siRNA seed was defined as the 2nd-7th position of the siRNA sequence. 2616 out of 3575 seeds occurred in four or more genes. On average, every seed occurred in 20.17 genes. If a seed associated with different genes was significantly enriched in the predicted genes list for a given GO term (Fisher test, p-value<0.01), then all the predictions linking the genes targeted by this seed to the enriched term were considered off-target effects and filtered out (Supplementary Fig. 5D).

### Subphenotypic space embedding

To generate the subphenotypic space that captures SVM decision boundaries we applied logistic regression to predict the SVM predictions (+/−) based on the selected features by the corresponding SVM term classifier. This resulted in approximate weights for the selected features by SVM. Then, t-SNE embedding was applied to the selected features by SVM term classifier multiplied by their corresponding weights.

### TCGA analysis

Data were obtained from the TCGA portal.

Silhouette clustering index of colorectal cancer molecular subtypes was used to determine clustering quality (separability and coherence of clusters) based on all genes or MSD genes. Principal component analysis was used to reduce the number of dimensions. The first 5 principal components were used to compute the silhouette index of colorectal cancer molecular subtypes as it resulted in the highest average silhouette value when all genes were considered. The same number of principal components was used when considering the top 300 predicted MSD genes based on SVM confidence. To estimate the significance of MSD genes in classification of colorectal cancer molecular subtypes, we generated a hundred random samples of 300 genes and computed the mean of average silhouette values for the hundred samples (t-test).

Transcription-based signatures: MSD signature was computed as the average expression of the top 300 MSD genes. Olfactory receptor signature was computed as the average expression of olfactory receptors associated with MSD. WNT signature was computed as the average of the following WNT genes: WNT1, WNT2, WNT2B, WNT3, WNT3A, WNT4, WNT5A, WNT5B, WNT6, WNT7A, WNT7B, WNT8A, WNT8B, WNT9A, WNT10A, and WNT10B. TGFβ signatures were computed as the average of genes identified in Calon *et al*(Calon *et al*, 2012)., except for endothelial-associated genes.

Expression state of olfactory receptors was empirically determined depending on the distribution of their expression values as following: OR51B4 cut-off = 0.25, OR5K1 cut-off = 0.01, and olfactory receptor metagene cut-off = 0.1. To compute clinically relevant olfactory receptor metagene for less expressed receptors, the expression state of each olfactory receptor was added iteratively to the metagene and retained if the metagene scored significant based on Kaplan Meier log-rank test (Supplementary Fig. 6E). 22 olfactory receptors were selected out of 84 genes.

Kaplan Meier survival analysis and Cox Proportional Hazard modelling were performed in R.

## Acknowledgments

We thank Dr. Victoria Green and Sophie Tritschler for generating the image-based dataset which provides a great resource for biological discovery. We also thank members of Pelkmans group, Christine Orengo and Jon Lees (UCL), Simon Leedham, Andrew Zisserman (University of Oxford) for useful discussions. HS is a Sir Henry Wellcome Fellow.

## Author contributions

HS conceived the work, designed and implemented KDML, performed all data analyses and wrote the paper. LP provided the image-based dataset and thoroughly discussed the results with HS. JR and LP reviewed and edited the paper.

## Supplementary Figure legends

**Supplementary Fig. 1.**
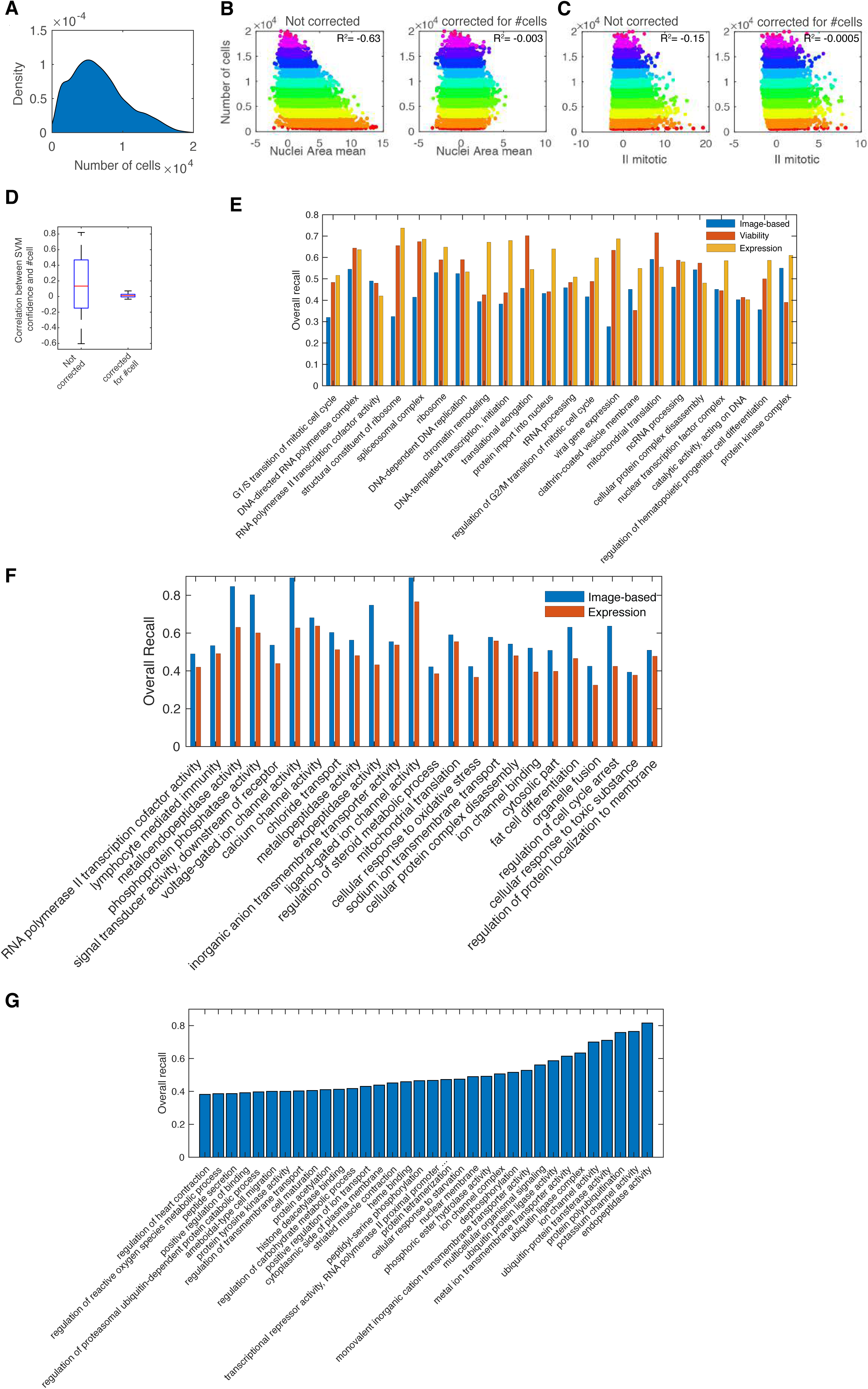
(A) Distribution of the number of cells across 18,033 gene knockdowns. (B-C) Scatter plots of TCN (total cell number) versus (B) Nuclei area mean and (C) II mitotic (infection index for mitotic cells) before and after correction for dependency on TCN. R^2^ indicates Pearson correlation coefficient. Colour indicates cell number (0-2000, … 18,000-20,000). (D) Box plots of the correlation between SVM confidence scores and cell number before and after TCN correction. Box plots elements: centre line, median; box limits, 25^th^ and 75^th^ percentiles; whiskers, +/−2.7 standard deviation. Points: outliers. (E) Overall recall of classifiable GO terms based on the three datasets. (F) GO terms where single cell-resolved image-based readouts outperform expression readouts based on overall recall. (G) GO terms that are only classifiable based on the image-based dataset.

**Supplementary Fig. 2.**
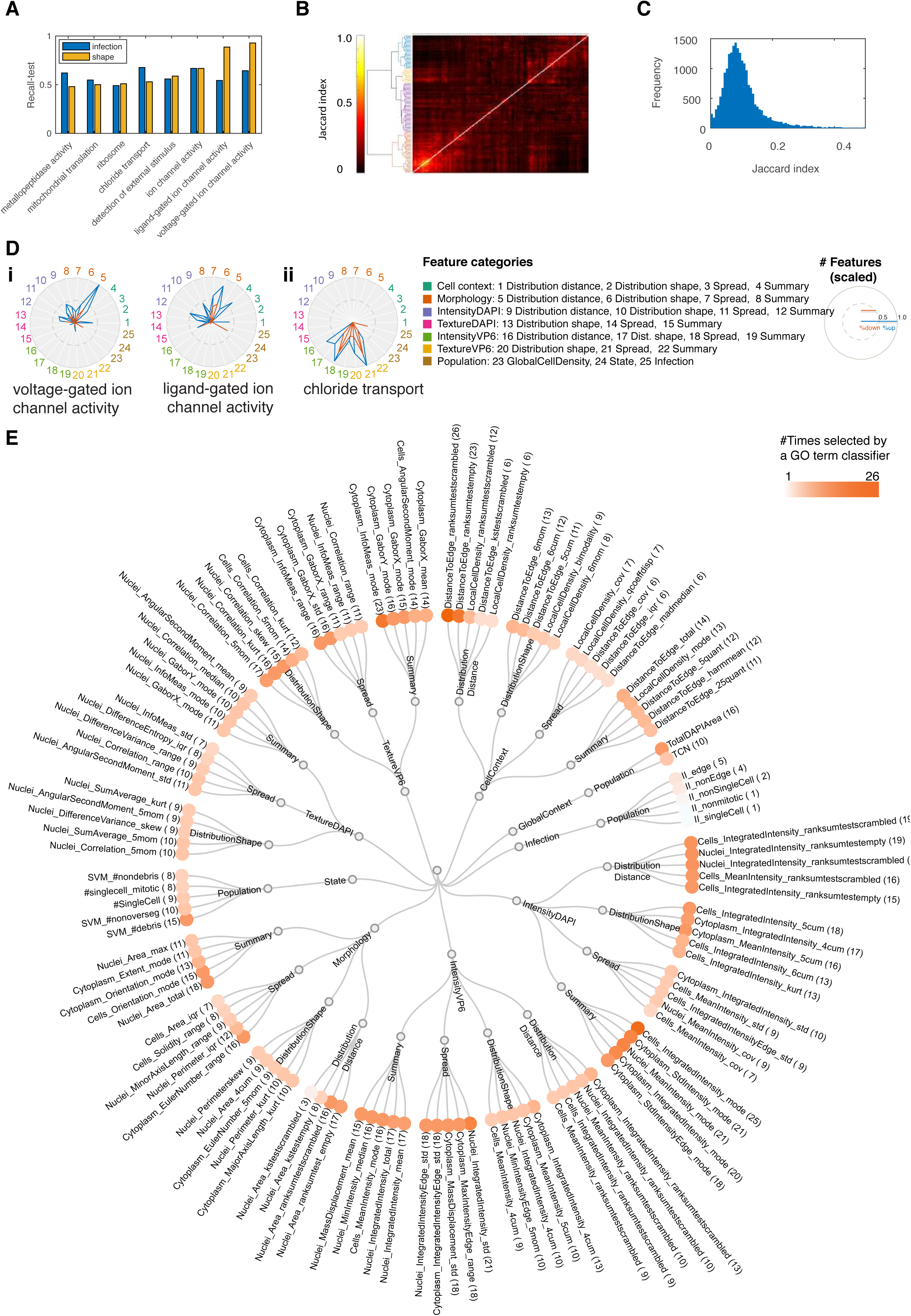
(A) The recall on test data for terms that are classifiable based on both infection and shape features. (B-C) Clustering (B) and distribution (C) of Jaccard index based on the overlap between predictions of GO term classifiers respectively. Most terms have a moderate overlap suggesting that different phenotypes are discovered. (D) The number of features in different categories that are selected by the respective GO term classifier (scaled) when (i) only shape features or (ii) only infection features are used for classification. Blue indicates the number of features with a higher average than control while red indicates the number of features with a lower average than control. (E) Feature categories based on feature type (i.e morphology, cell context, DAPI intensity), measurement type (i.e. summary, spread, distribution shape). The top most frequently selected features by different GO term classifiers are shown as examples if applicable. Hue of orange circles indicates feature importance based on the number of term classifiers selecting the corresponding feature.

**Supplementary Fig. 3.**
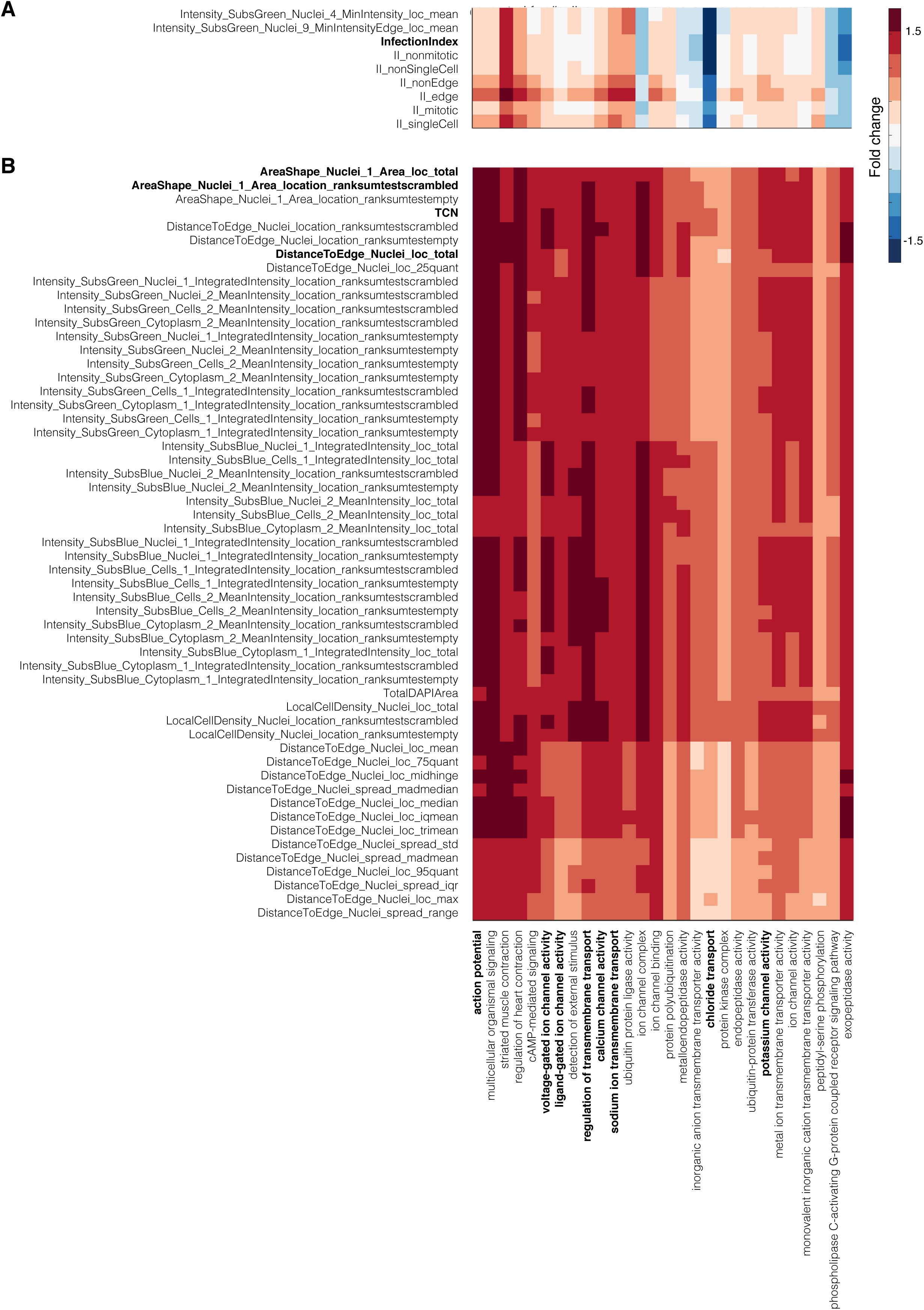
(A-B) Examples on significantly changed features for terms in Fig. 3A (C1) which include many membrane transport terms. TCN: total cell number.

**Supplementary Figure 4.**
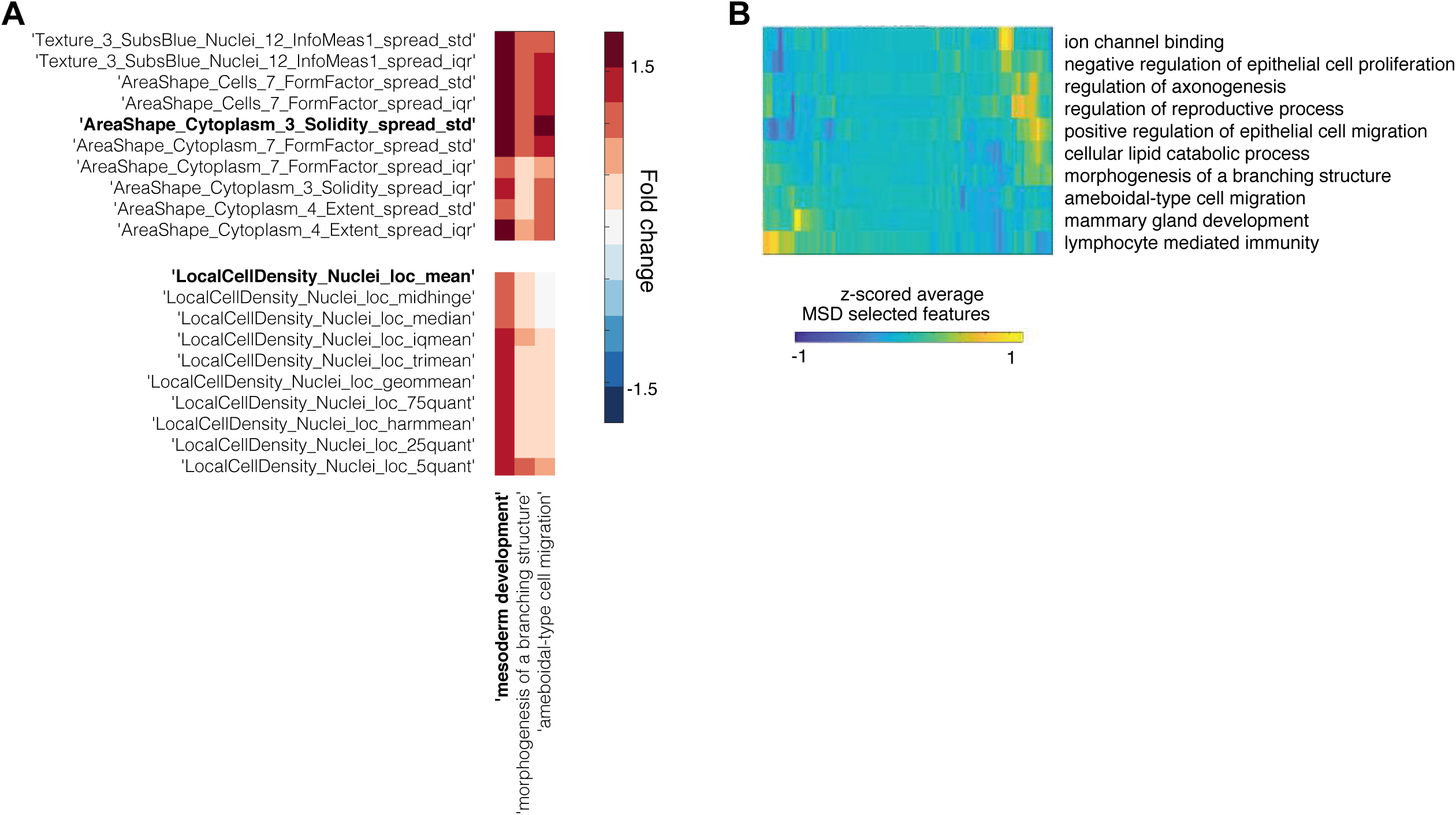
(A) Example of most significantly changed features for terms in Fig. 3A (C10) including MSD. (B) Average values of features selected by MSD classifier for genes predicted to perform additional functions other than MSD.

**Supplementary Figure 5.**
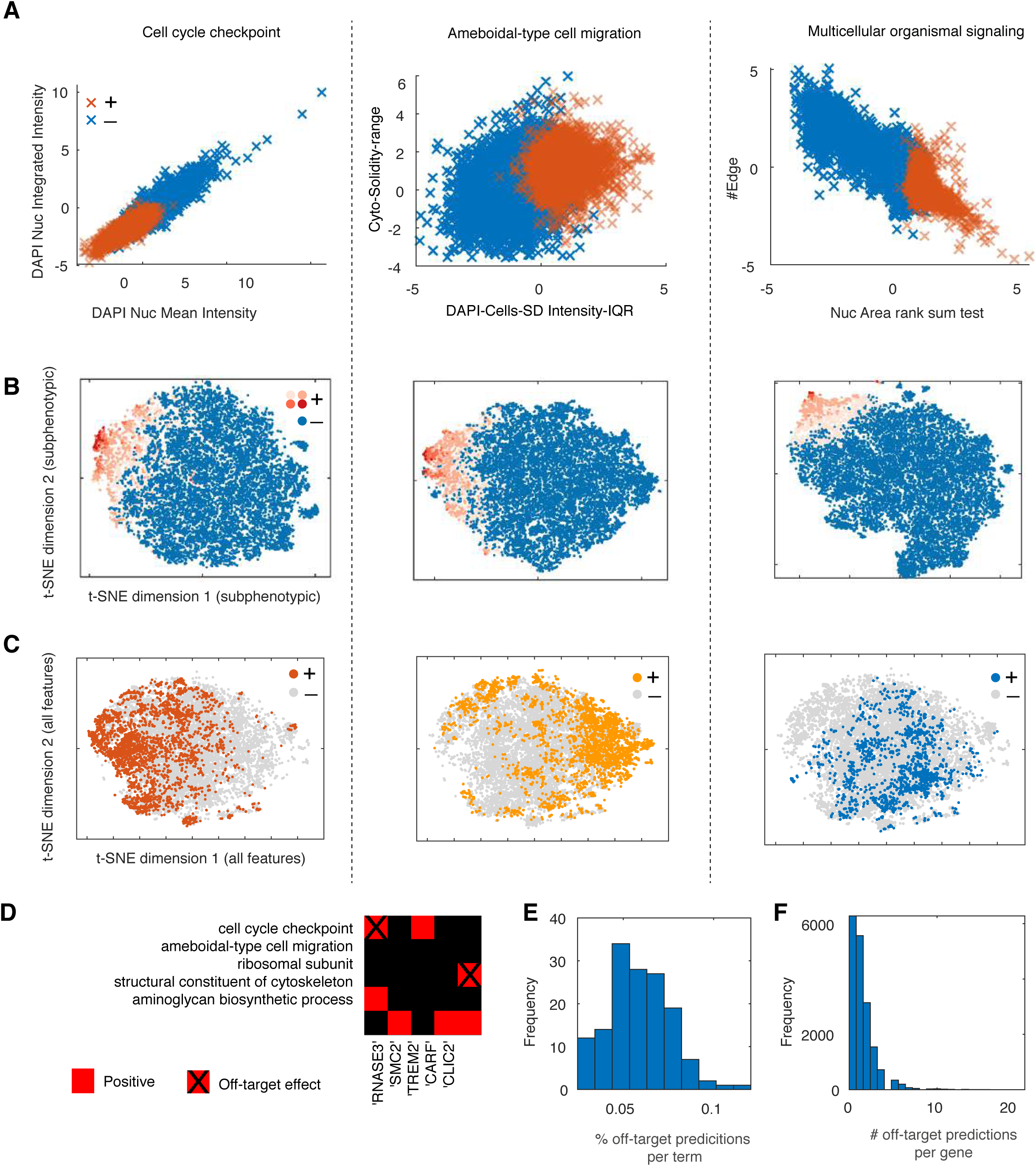
(A) Comparison between negative and positive genes predicted by the respective GO term classifier based on (a) two significantly changed feature, (B) subphenotypic t-SNE embedding (based on the selected features by the respective classifier) where red hues indicate SVM rank, or (C) phenotypic t-SNE embedding (based on all measured features). (D) Example on predicted functions for five genes and the detected off-target effects based on siRNA seed enrichment. (E-F) Distribution of the number of off-target predictions per GO term classifier (E) or per gene (F).

**Supplementary Figure 6.**
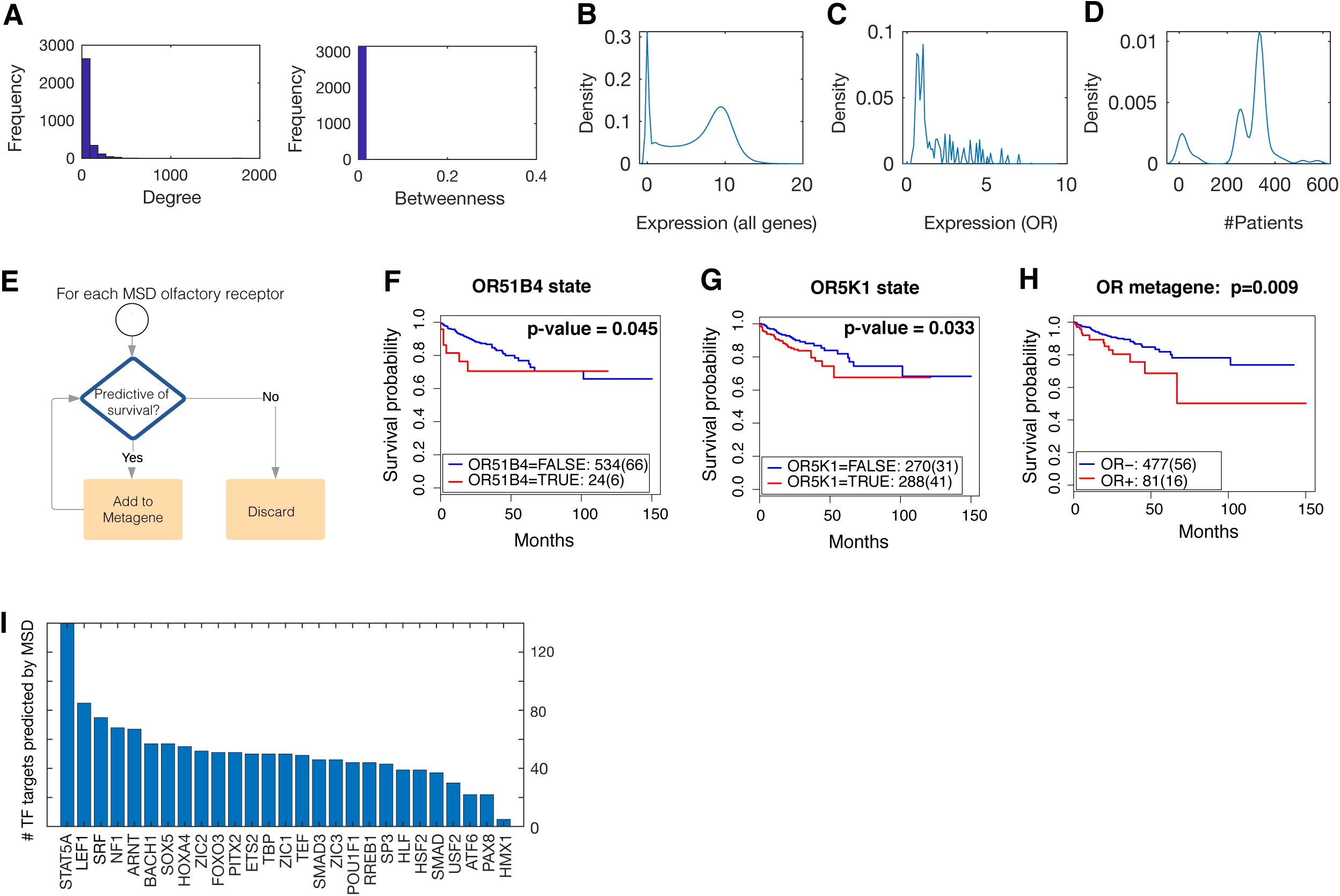
(A) Distribution of network degree and betweenness centrality measures for the network in Fig. 6B show a lack of gene hubs. (B-C) Distribution of the expression of all genes (B) and MSD-associated olfactory receptors (C) in colorectal cancer patients based on TCGA. (D) Distribution of the number of patients where an MSD-associated olfactory receptor is expressed. (E-F) Kaplan Meier survival analysis of colorectal cancer patients based on the expression state of *OR51B4* (F), *OR5K1*+ (G) and MSD-associated olfactory receptor metagene (H). OR: olfactory receptor. (I) Derivation of MSD-associated olfactory receptor metagene where the expression state of each receptor is added iteratively to the metagene and retained if scored significant based on Kaplan Meier survival test. (K) Wald statistic value based on Cox Proportional Hazard analysis of the predictivity of olfactory receptors metagene of survival against tumour grade, presence of lymph nodes and metastasis. * indicates p-value<0.05 and ** indicates p-value <0.001. Detailed results for all tested variables and model significance are shown in Supplementary Table 7.

## Supplementary Tables

**Supplementary Table 1:** list of included GO terms and their classifiability based on the different datasets.

**Supplementary Table 2:** list of aggregated features of single cell image-based data and their category assignment.

**Supplementary Table 3:** list of high confidence predictions based on top overlapping terms between expression, viability and image-based datasets.

**Supplementary Table 4:** List of MSD genes in overrepresented KEGG pathways.

**Supplementary Table 5:** Highest confidence predictions based on image-based morphology and infections features.

**Supplementary Table 6:** list of MSD associated olfactory receptors and members of the olfactory receptor metagene.

**Supplementary Table 7-9:** Results of Cox Proportional Hazard model for OR51B4, OR5K1, and olfactory receptor metagene respectively.

